# Sex differences in agonistic social behavior: Effects of exercise and ovariectomy

**DOI:** 10.64898/2026.11.05.764709

**Authors:** Jessica D. Westerman, Margaret K. Tanner, Alyssa A Hohorst, Lilia Khellaf, Filsan H. Mawlid, Ciana M. Beller, Benjamin N. Greenwood

## Abstract

Aggressive behavior encompasses a range of offensive and defensive responses that may be differentially influenced by sex, ovarian hormones, and physical activity. However, much of the existing research has focused on males and utilized paradigms designed to elicit aggression, leaving the expression and regulation of offensive and defensive behaviors during other forms of social interaction poorly understood. Here, we investigated the effects of sex, voluntary wheel running (VWR), and ovariectomy (OVX) on agonistic behavior during juvenile social exploration in adult Sprague-Dawley rats. Males exhibited greater offensive aggression, characterized by increased aggressive grooming and pinning, whereas females displayed more defensive kicking, darting, and avoidance. OVX selectively reduced defensive kicking without significantly affecting offensive aggression or avoidance, suggesting that distinct behavioral responses differ in their sensitivity to ovarian hormone removal. In contrast, VWR did not significantly alter any of the behaviors examined. Importantly, exploratory composite analyses revealed a significant sex difference in offensive aggression that was no longer detected when defensive behaviors were incorporated into the measure. Together, these findings demonstrate that sex differences in agonistic behavior extend beyond overall levels of aggression to distinct patterns of offensive and defensive responses. Distinguishing these behavioral categories may therefore be important for identifying how various manipulations, such as sex, ovarian hormones, and physical activity, influence agonistic behavior.

## Introduction

Aggressive behavior represents a significant public health concern. In 2024, an estimated 6.7 million violent victimizations occurred among individuals aged 12 years and older in the United States alone (Tapp & Coen, 2025). Between 2019 and 2021, assault-related emergency department visits were highest among individuals aged 18–24, with an average annual rate of 9.2 visits per 1,000 people (Davis & Santo, 2023). Notably, sex differences in violence-related outcomes vary with injury severity. Between 2015 and 2020, males accounted for approximately 86% of homicides among individuals aged 10–24, compared with 61% of nonfatal assault-related emergency department visits (Peterson et al., 2023). Given the prevalence and consequences of aggressive behavior, identifying factors that may reduce its expression is of high importance.

Physical activity has emerged as a potential factor to reduce aggressive behavior, with studies in humans reporting associations between greater physical activity and reduced aggression (Yue et al., 2025). A recent meta-analysis further found that physical activity reduced self- or observer-rated aggression in children and adolescents, with significant reductions in physical aggression, anger, and hostility, but not verbal aggression (Zhao & Wang, 2025). Another meta-analysis found that physical activity interventions significantly reduced aggression rated by others, but not self-reported aggression, and no significant effects on aggression subtypes, such as physical, verbal, and indirect aggression, were noted (Ouyang & Liu, 2023). Mixed findings have also been reported in rodents. In male mice, voluntary wheel running (VWR) has been shown to reverse social-isolation-induced increases in aggression during resident-intruder testing (Ren & Tai, 2023), whereas other work has found that VWR did not significantly reduce the number of attacks in socially isolated male mice (Umamichi et al., 2026).

Conflicting results on the effects of exercise on aggression could be due in part to a narrow definition of aggressive behavior (e.g., “physical aggression” in humans, “aggression” and “attacks” in rodents), rather than distinguishing between different forms. Aggressive behavior in rodents, for example, varies with the circumstances of the social interaction and encompasses a range of responses that can be broadly distinguished as offensive or defensive. Offensive aggression is typically associated with the initiation of agonistic interactions, whereas defensive behaviors occur in response to perceived threats or unwanted social encounters (Blanchard & Blanchard, 1989; Koolhaas et al., 2013). These behavioral categories can be differentially influenced by neurobiological and pharmacological manipulations, suggesting that they are regulated, at least in part, by distinct mechanisms (Blanchard & Blanchard, 1989). Distinguishing offensive from defensive responses may therefore reveal differences that are obscured when agonistic behavior is treated as a single outcome. Understanding the effects of physical activity on aggressive behavior in rodents is particularly important, as rodent models can be used to identify underlying mechanisms and potentially novel biological targets for the reduction of aggression.

Research on aggression has historically focused predominantly on males (Been et al., 2019), a pattern also reflected in studies investigating the effects of exercise on aggressive behavior (Hoffmann et al., 1987; Ren & Tai, 2023). The limited existing studies that include both sexes have revealed some differences in aggressive behavior between sexes. While some studies have concluded that males generally exhibit higher levels of overt physical aggression than females across rodent species (Bell, 2018; Been et al., 2019), sex differences in aggression extend beyond overall levels of aggressive behavior to differences in its regulation and expression. For example, studies in Syrian hamsters have identified sex differences in the neurochemical regulation of offensive aggression, characterized by behaviors such as attacking and biting. Specifically, hypothalamic arginine vasopressin has been reported to facilitate offensive aggression in males but inhibit it in females (Terranova et al., 2017). Work in mice has also identified sex differences in behavioral expression despite similar overall levels of aggression: males displayed more persistent attack sequences, whereas females exhibited more intermittent aggression interspersed with investigative behaviors (Aubry et al., 2022). These findings suggest that sex differences in aggression may be reflected in the specific behaviors expressed. However, because many studies have used paradigms designed to elicit aggression, such as the resident-intruder test (Koolhaas et al., 2013; Miczek, 1979; Aubry et al., 2022; Takahashi et al., 2014), it remains unclear whether these differences extend to offensive and defensive responses during social interactions not explicitly designed to provoke aggression.

In addition to the social context in which aggression is assessed, ovarian hormone status may influence the expression of agonistic behavior in females, although previous studies have reported inconsistent findings. Ovariectomy has been reported to leave aggression unchanged under some experimental conditions (DeBold & Miczek, 1984) and to reduce it under others (Ho et al., 2001). Aggressive behavior has also been reported to vary across the estrous cycle, with females exhibiting greater aggression during nonreceptive phases than during receptive phases (Ho et al., 2001). These findings suggest that the relationship between ovarian hormones and aggression may depend on the experimental context and hormonal state, raising the question of whether offensive and defensive responses differ in their sensitivity to ovarian hormone removal.

Although social encounters between adult rodents are often aggressive, interactions between adult and juvenile rats are predominantly non-aggressive (Blanchard & Blanchard, 2003). Although minimal compared to aggressive interactions between adults, we have observed anecdotally in prior studies that aggression in adults towards juveniles can occur. Juvenile social interaction thus provides an opportunity to examine agonistic behaviors in adult rats during social interactions not specifically designed to provoke aggression. In the present study, we investigated the effects of prior VWR on offensive and defensive aggressive behaviors in male and female Sprague-Dawley rats under conditions that do not provoke overt aggression. Offensive behavior was classified as aggressive grooming, pinning, and attacking, while darting, kicking, and avoidance were scored as defensive behaviors. In addition to investigating sex differences in the occurrences of these behaviors and the effects of exercise, the contribution of ovarian hormones to female-specific behavioral patterns were assessed by comparing ovariectomized and sham-operated females. We hypothesized that exercise would reduce aggression in both sexes, with potential sex-dependent patterns of aggression being dependent on the presence of ovarian hormones.

We found a sex-specific pattern of agonistic behavior, whereby males displayed more offensive aggression than females, but females displayed more defensive aggression and avoidance than males. Interestingly, the female-specific pattern of defensive aggression was dependent on the presence of ovarian hormones. Exercise did not significantly impact any of the behaviors measured. Together, results highlight the importance of distinguishing offensive and defensive responses when examining factors influencing the expression of agonistic behavior.

## Methods

### Animals and housing

The present study utilized a cohort of adult, age-matched, female (n= 76) and male (n= 22) Sprague Dawley rats (Envigo, Indianapolis, IN, USA) previously described in Tanner et al. (2023). The prior publication examined sex differences in the effects of VWR on stress-induced anxiety-like behavior (Tanner et al., 2023), whereas the present analysis focuses on sex differences in the effects of VWR on aggression that have not been previously reported. Although the original publication included rats exposed to an inescapable stressor, the current analysis focused on sex differences in effects of VWR on agonistic behavior by analyzing only the non-stressed animals. Rats were approximately 7 weeks old at time of arrival and were single-housed in Nalgene Plexiglas cages (45.5 × 24 × 21 cm, length × width × height). Rats were housed with locked wheels (1.081 m circumference; Starr Life Sciences, Oakmont, PA) upon arrival, and wheels in the cages of VWR were unlocked after 1 week of acclimation to the vivarium. Rats were singly housed to permit individual measurement of VWR activity. Previous work has demonstrated that single versus pair housing does not significantly alter the effects of VWR (Greenwood & Fleshner, 2011). Food (Teklad 2020X rodent diet, Envigo) and water were available *ad libitum*. Body weights were recorded weekly. Rats were maintained on a 12:12 h light-dark cycle (lights on 0600–1800 h) in a temperature-controlled (22 °C) and humidity-controlled (∼30% relative humidity) vivarium accredited by the Association for Assessment and Accreditation of Laboratory Animal Care International (AAALAC). All experimental procedures were approved by the Institutional Animal Care and Use Committees at the University of Colorado Denver.

### Ovariectomy

Surgical procedures have been described previously (Tanner et al., 2023). Female rats underwent either a bilateral ovariectomy (OVX; n = 28) or SHAM surgery (SHAM; n = 21). Rats were anesthetized with ketamine (75 mg/kg, i.p.) and medetomidine (0.5 mg/kg, i.p.) with atipamezole hydrochloride (Antisedan, 2mg/kg, i.p.) used as a reversing agent. A single dorsal incision was made, followed by bilateral incisions through the underlying muscle wall. The ovaries were exteriorized, the uterine horns were ligated using absorbable Vicryl suture (4-0, FS-2), and the ovaries were removed. Muscle and skin incisions were subsequently closed with absorbable sutures. Bupivacaine (2 mg/kg) was applied to the muscle incisions prior to closure. SHAM surgeries followed the same procedure, except that the ovaries were left intact. Carprofen (5□mg/kg s.c.) and penicillin G (22,000□IU/rat s.c.) were administered at induction and every 24□h for 72□h after surgery. Rats were allowed to recover for 2□weeks before further experimental procedures. Estrous cycle was monitored after surgery to verify SHAM-operated rats were displaying normal estrous cyclicity, and OVX rats were not displaying vaginal cytology consistent with proestrus or estrus. No rats were removed from the study due to failure of SHAM or OVX defined by these criteria (Tanner et al., 2023).

### Voluntary Wheel Running

To compare the effects of VWR on aggressive behavior between sexes, male and female rats were randomly assigned to either 3- or 6-week VWR (VWR; n = 20 females, 14 males) or sedentary (Sed; n = 7 females, 8 males) conditions. These durations were used in the original publication to investigate whether there are sex differences in the duration of VWR required for stress resistance (Tanner et al., 2023). To determine whether exercise duration influenced aggressive behavior, a two-way ANOVA was performed with sex (male vs. female) and treatment group (sedentary, 3-week exercise, and 6-week exercise) as factors to assess differences in agonistic behaviors based on duration spent running. There were no significant sex × duration interactions observed for aggressive grooming (*F*(2, 43) = 3.10, *p* = .0551), darting (*F*(2, 43) = 1.47, *p* = .2424), pinning (*F*(2, 43) = 0.21, *p* = .8089), kicking (*F*(2, 43) = 0.07, *p* = .9329), attacking (*F*(2, 43) = 0.97, *p* = .3875), or avoidance (*F*(2, 43) = 0.48, *p* = .6192); thus, 3 and 6 week exercise animals were combined into one VWR group for each sex.

Unequal sample sizes between sexes reflected the original experimental design, which included additional females to investigate the influence of estrous cycle phase and a separate ovariectomy experiment. A total of 11 animals were excluded from the sex × exercise analysis because juvenile social exploration recordings were missing or corrupted (female sedentary, n = 5; female VWR, n = 4; male VWR, n = 2).

To assess the effects of VWR and OVX on defensive behaviors in females, rats were assigned to either 3 weeks of VWR (OVX, n = 15; SHAM, n = 12) or sedentary conditions (OVX, n = 13; SHAM, n = 9) following OVX or SHAM surgery. For rats assigned to VWR, running wheels remained unlocked throughout the exercise period, whereas wheels remained immobilized for sedentary rats. The running wheel was the only form of cage enrichment provided.

Running activity was automatically recorded every minute using VitalView software (Starr Life Sciences). Running data have been published previously and included running speed, number of running bouts, distance per bout, and bout duration (Tanner et al., 2023).

### Juvenile Social Exploration

Juvenile social exploration was used to assess behavioral interactions between adult experimental rats and a juvenile rat of the same sex. Juvenile social exploration testing occurred 24 h after the end of the VWR or sedentary condition. Experimental rats were individually transferred to a clean Nalgene Plexiglas cage with bedding in a brightly lit testing room and allowed to habituate for one hour. Following habituation, a juvenile rat (age P28 ± 2 d) of the same sex and species was placed in the same cage as the experimental rat.

Interactions were recorded for three minutes, and behavior was later scored with a custom program from recorded videos by two lab personnel blind to treatment conditions. Final scores represent an average of the scores from the 2 experimenters, whose inter-rater reliability was assessed using a two-way, absolute-agreement, single-measure intraclass correlation coefficient (ICC). Inter-rater reliability was excellent across all six behavioral measures (mean ICC = 0.961; range = 0.933–0.985; n = 98). Rather than quantifying total social investigation as occurred in the prior publication (Tanner et al., 2023), behavioral scoring focused on offensive aggression and defensive responses displayed by the adult experimental rat. Scored behaviors classified as offensive included aggressive grooming, pinning, and attacking, whereas kicking was classified as defensive aggression. Darting and avoidance were classified as nonaggressive defensive responses. Behaviors were scored according to predefined operational criteria. Aggressive grooming was defined as forceful grooming of the juvenile by the adult, typically involving restraint or persistent physical contact. Attacking was defined as forceful restraint accompanied by biting directed toward the juvenile. Pinning was defined as the adult physically restraining the juvenile against the cage floor using the forepaws and/or a hindlimb. Kicking was defined as forceful contact used to repel the juvenile. Darting was defined as rapid, sudden movement by the adult across the cage that may reflect disengagement or escape behavior. Avoidance was defined as active disengagement, withdrawal, or movement away from the juvenile.

### Statistical Analyses

Behavioral scores from two independent raters were averaged for each animal before statistical analysis. Inter-rater reliability was assessed using a two-way, absolute-agreement, single-measure intraclass correlation coefficient (ICC). The effects of sex and VWR on offensive and defensive behaviors were analyzed using two-way ANOVAs with sex and exercise as between-subjects factors. In the ovariectomy experiment, two-way ANOVAs were conducted with OVX status and exercise as between-subjects factors. Type III sums of squares were used for all ANOVAs. Assumptions of normality and homogeneity of variance were evaluated using Shapiro–Wilk tests of model residuals and Levene’s tests, respectively. Shapiro–Wilk tests indicated significant departures from normality for all behavioral outcomes, while Levene’s tests identified unequal variances for some outcomes. Therefore, all ANOVAs were conducted using heteroscedasticity-consistent (HC3) robust standard errors. Where appropriate, post hoc pairwise comparisons of estimated marginal means were performed using HC3-robust standard errors with Holm’s correction for multiple comparisons. For exploratory composite analyses, behavioral measures were standardized across all animals using z-scores, calculated by subtracting the sample mean from each individual value and dividing by the sample standard deviation. Standardized scores were then summed to generate three composite measures: offensive aggression (aggressive grooming, pinning, and attacking), physical aggression (offensive behaviors plus defensive kicking), and broader agonistic behavior (physical aggression plus darting). This approach allowed behaviors measured in different units (durations and frequencies) to contribute equally to each composite. Statistical significance was defined as p < 0.05. Data are presented as mean ± SEM.

## Results

### Males display more offensive aggressive behavior than females, and exercise does not influence offensive aggression

Here, we assessed the effects of sex and exercise on offensive aggressive behaviors during juvenile social exploration. Females displayed less aggressive grooming (main effect of sex: *F*(1, 45) = 8.13, *p* = 0.0065, Figure 1A) and pinning (main effect of sex: *F*(1, 45) = 5.94, *p* = 0.0189; Figure 1B) than males, whereas the number of attacks did not differ between sexes (main effect of sex: *F*(1, 45) = 0.020, *p* = 0.8870; Figure 1C). There was a trend for VWR to reduce aggressive grooming (main effect of exercise: *F*(1, 45) = 3.47, *p* = 0.0689; sex × exercise interaction: *F*(1, 45) = 2.87, *p* = 0.0972; Figure 1A), but VWR did not impact pinning (main effect of exercise: *F*(1, 45) = 0.007, *p* = 0.9331; sex × exercise interaction: *F*(1, 45) = 0.13, *p* = 0.7200; Figure 1B), or attacks (main effect of exercise: *F*(1, 45) = 0.94, *p* = 0.3362; sex × exercise interaction: *F*(1, 45) = 2.23, *p* = 0.1420; Figure 1C).

**Figure 1.**
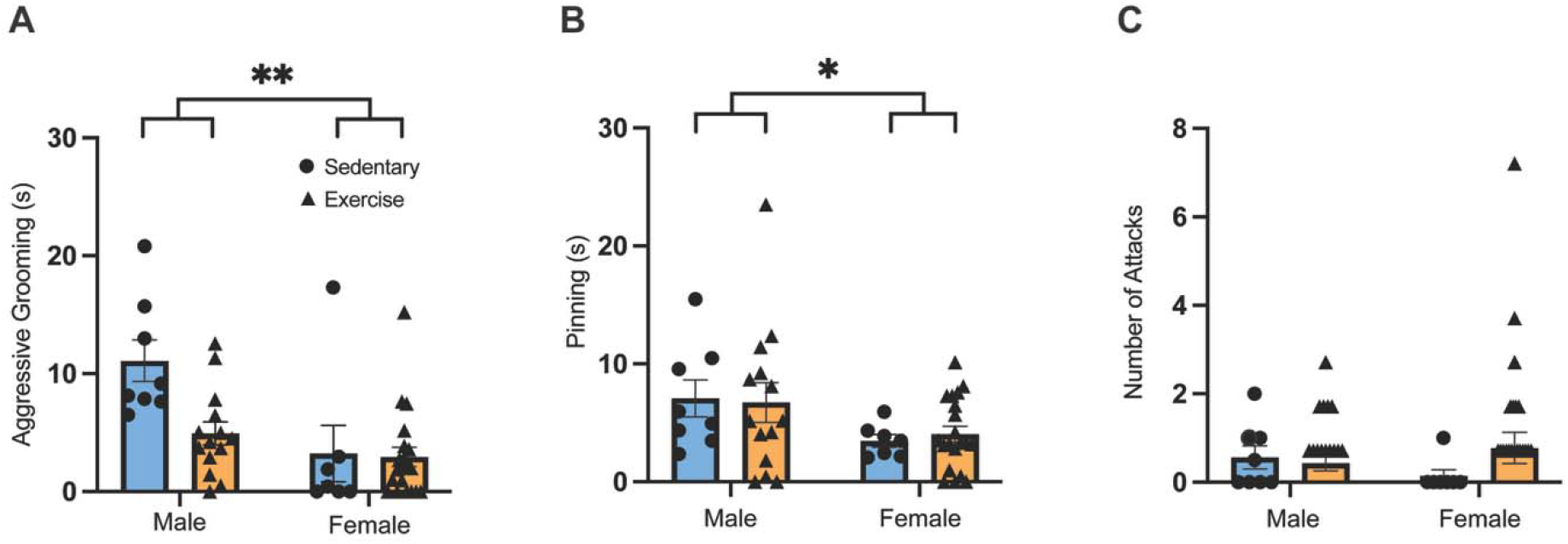
Effects of exercise and sex on offensive aggressive behaviors. Aggressive behaviors were quantified in female sedentary (*n* = 7), female exercise (*n* = 20), male sedentary (*n* = 8), and male exercise (*n* = 14) rats. **(A)** Aggressive grooming, **(B)** pinning, and **(C)** number of attacks. Post hoc pairwise comparisons of estimated marginal means were performed with Holm’s correction for multiple comparisons. \**p* < .05, \*\**p* < .01. Bars represent group means ± SEM; circles or triangles represent individual animals.

### Exercise does not impact female-specific defensive behavior

We next analyzed the effects of sex and exercise on defensive behaviors during juvenile social exploration. Females displayed more kicking behavior than males regardless of exercise condition (main effect of sex: *F*(1, 45) = 19.03, *p* < 0.0001; Figure 2A). Females also displayed significantly more darting than males (main effect of sex: *F*(1, 45) = 5.12, *p* = 0.0285; Figure 2B). Finally, females also spent more time avoiding the juvenile than males (main effect of sex: *F*(1, 45) = 26.69, *p* < 0.0001; Figure 2C). VWR had no significant effect on kicking (main effect of exercise: F(1, 45) = 0.13, p = 0.7209; sex × exercise interaction: F(1, 45) = 0.19, p = 0.6613; Figure 2A), darting (main effect of exercise: F(1, 45) = 0.020, p = 0.8874; sex × exercise interaction: F(1, 45) = 0.52, p = 0.4765; Figure 2B), or avoidance (main effect of exercise: F(1, 45) = 0.029, p = 0.8645; sex × exercise interaction: F(1, 45) = 0.038, p = 0.8457; Figure 2C).

**Figure 2.**
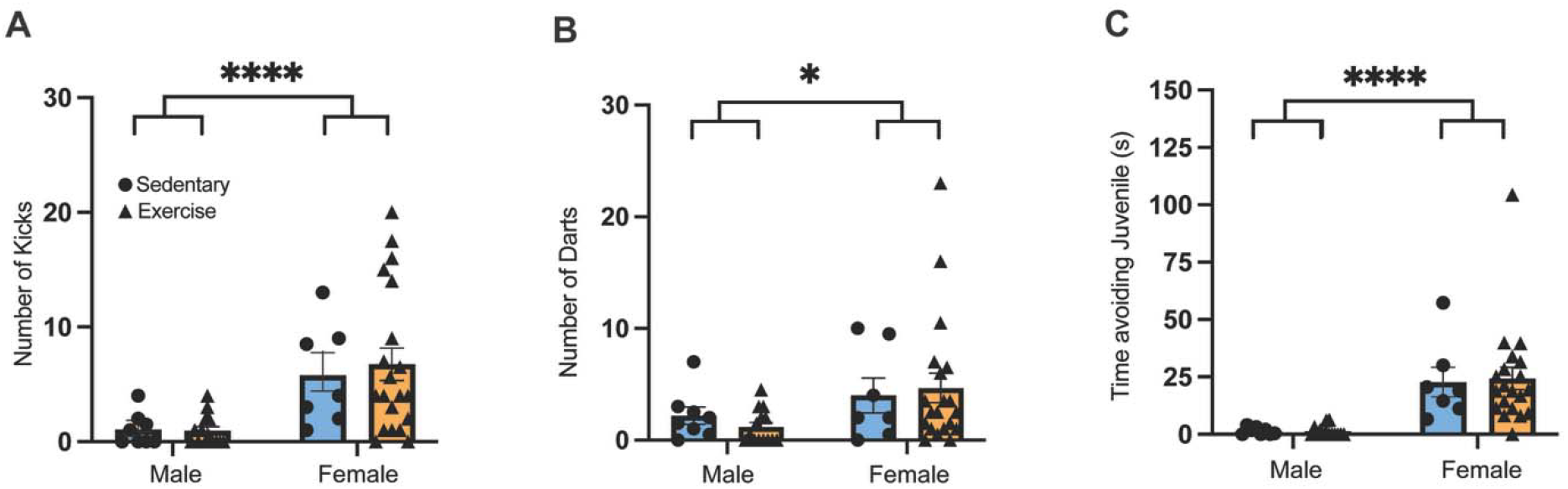
Effects of sex and exercise on defensive behavior. Defensive behaviors were quantified in female sedentary (*n* = 7), female exercise (*n* = 20), male sedentary (*n* = 8), and male exercise (*n* = 14) rats. **(A**) Kicking, **(B)** darting, and **(C)** time spent avoiding the juvenile. \**p*<.05 \*\**p* < .001, \*\*\*\**p* < .0001. Bars represent mean ± SEM; circles or triangles represent individual animals.

### Ovariectomy does not impact offensive aggression

We next evaluated whether ovarian hormone reduction altered the expression of offensive aggressive behaviors in female rats. Exercise again did not impact offensive aggressive behaviors in females, regardless of hormone status. Neither aggressive grooming (main effect of exercise: *F*(1, 45) = 0.20, *p* = 0.6559; Figure 3A), pinning (main effect of exercise: *F*(1, 45) = 0.26, *p* = 0.6133; Figure 3B), nor number of attacks (main effect of exercise: *F*(1, 45) = 1.74, *p* = 0.1938; Figure 3C) were significantly affected by exercise. OVX also did not alter aggressive grooming (main effect of OVX: *F*(1, 45) = 1.86, *p* = 0.1791; OVX × exercise interaction: *F*(1, 45) = 0.003, *p* = 0.9558; Figure 3A), pinning (main effect of OVX: *F*(1, 45) = 0.049, *p* = 0.8253; OVX × exercise interaction: *F*(1, 45) = 0.92, *p* = 0.3431; Figure 3B), or number of attacks (main effect of OVX: *F*(1, 45) = 0.11, *p* = 0.7438; OVX × exercise interaction: *F*(1, 45) = 1.07, *p* = 0.3071; Figure 3C).

**Figure 3.**
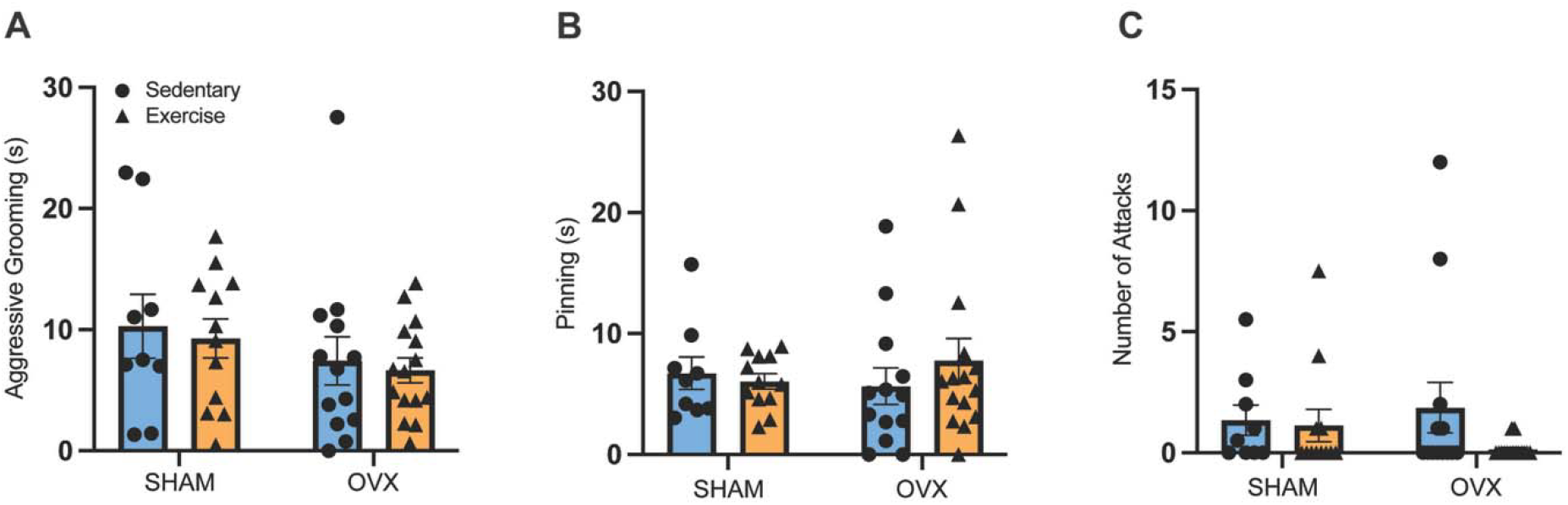
Effects of ovariectomy and exercise on offensive aggression behaviors. Aggressive behaviors were quantified in OVX sedentary (*n* = 13), OVX exercise (*n* = 15), SHAM sedentary (*n* = 9), and SHAM exercise (*n* = 12) rats. **(A)** Aggressive grooming, **(B)** pinning, and **(C)** number of attacks. Bars represent mean ± SEM; individual circles and triangles represent individual animals.

### Ovariectomy reduces defensive kicking but not avoidance behaviors

OVX status differentially impacted defensive behaviors. There was a main effect of OVX status on kicking behavior, with OVX females displaying fewer kicks than SHAM females ((*F*(1, 45) = 12.29, *p* = 0.0010; Figure 4A). Neither darting (*F*(1, 45) = 3.35, *p* = 0.0739; Figure 4B) nor time spent avoiding the juvenile (*F*(1, 45) = 1.16, *p* = 0.2880; Figure 4C) were significantly impacted by OVX. Exercise did not significantly affect any of the defensive behaviors examined (darting: *F*(1, 45) = 1.22, *p* = 0.2761; kicking: *F*(1, 45) = 0.16, *p* = 0.6938; avoidance: *F*(1, 45) = 1.77, *p* = 0.1906), and no significant OVX × exercise interactions were observed (darting: *F*(1, 45) = 0.91, *p* = 0.3453; kicking: *F*(1, 45) = 0.27, *p* = 0.6083; avoidance: *F*(1, 45) = 1.08, *p* = 0.3053).

**Figure 4.**
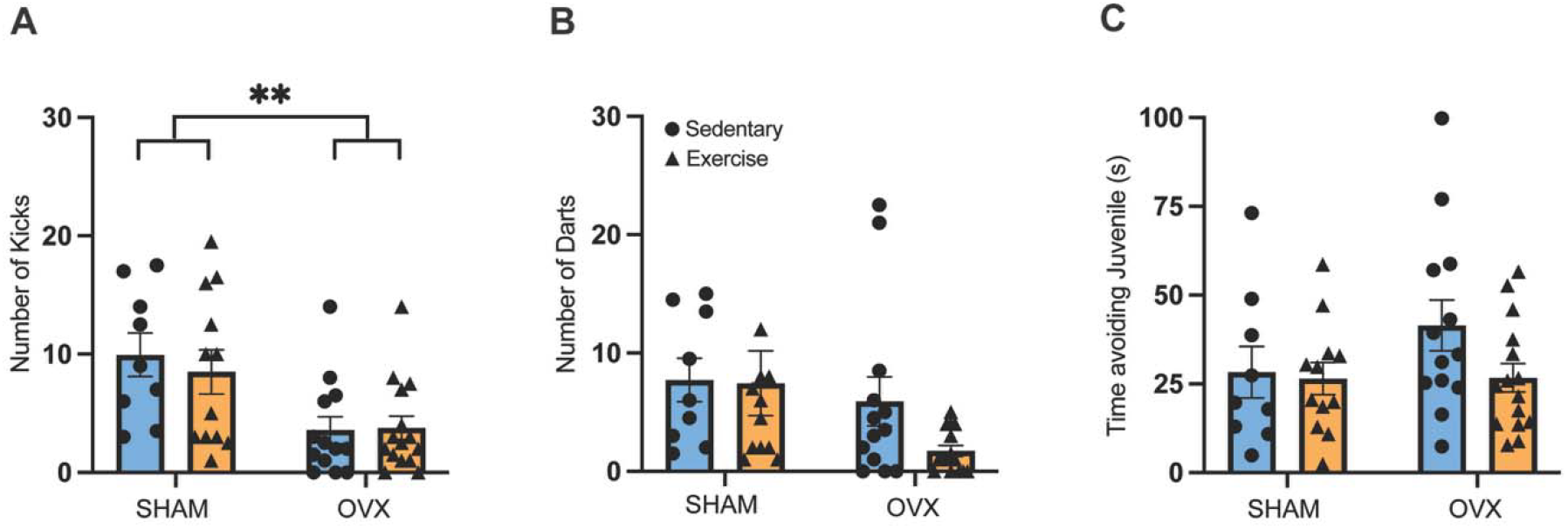
Effects of ovariectomy and exercise on defensive and avoidance behaviors. Behaviors were quantified in OVX sedentary (*n* = 13), OVX exercise (*n* = 15), SHAM sedentary (*n* = 9), and SHAM exercise (*n* = 12) rats. **(A)** Kicking, **(B)** darting, and **(C)** time spent avoiding the juvenile. Bars represent mean ± SEM; individual circles and triangles represent individual animals. \*\**p* < .01.

### Inclusion of defensive behaviors eliminates significant sex differences in composite aggression scores

To determine whether the inclusion of defensive behaviors influences the detection of sex differences in agonistic behavior, composite scores were calculated from standardized behavioral measures (Figure 5). A significant main effect of sex was observed for the offensive aggression composite, which included aggressive grooming, pinning, and attacking, with males exhibiting higher scores than females (F(1, 45) = 8.49, p = .0055; Figure 5A). However, this sex difference was no longer significant when defensive kicking was included in the composite (F(1, 45) = 0.89, p = .3511; Figure 5B) or when darting was additionally incorporated (F(1, 45) = 0.008, p = .9274; Figure 5C). Neither exercise nor the sex × exercise interaction significantly affected any composite score (all p > .05). These findings indicate that the detection of sex differences in agonistic behavior depends on whether defensive behaviors are included in the composite measure.

**Figure 5.**
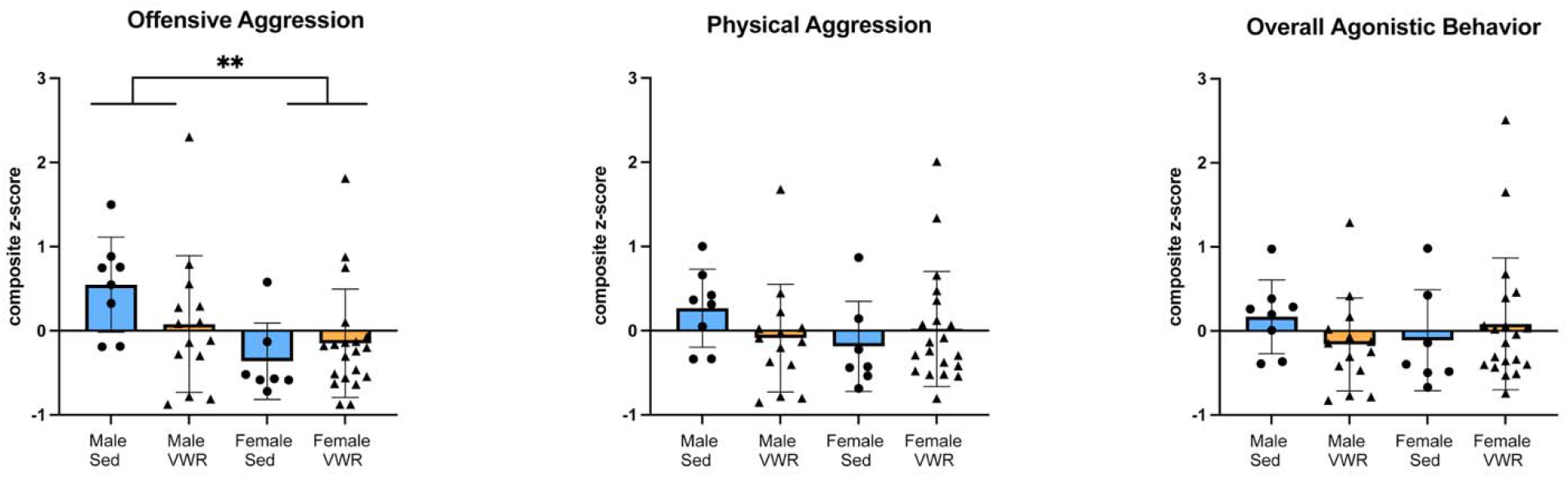
Inclusion of defensive behaviors alters the detection of sex differences in agonistic behavior. Composite scores were calculated from standardized behavioral measures during juvenile social exploration. (A) The offensive aggression composite including aggressive grooming, pinning, and attacking (B) The physical aggression composite additionally including defensive kicking. (C) The agonistic composite further incorporates darting. Bars represent mean ± SEM, with individual points representing individual rats. **p < .01, main effect of sex.

## Discussion

The present study investigated effects of VWR, sex, and ovarian hormone status on offensive and defensive behavior displayed during juvenile social exploration. As a social paradigm, juvenile social exploration is not designed to elicit overt aggression, thus allowing examination of agonistic behaviors that occur spontaneously during social interaction, rather than in response to an explicitly aggressive challenge. Although minimal, adult aggression toward the juvenile did occur. Males exhibited greater offensive aggression, as reflected by increased aggressive grooming and pinning, whereas females displayed more defensive kicking and avoidance behaviors. VWR did not significantly affect any of the aggressive or defensive behaviors examined, although a nonsignificant trend toward reduced aggressive grooming in males was observed. Ovariectomy selectively reduced the female pattern of defensive aggression (kicking), while not significantly altering offensive aggression or avoidance behaviors. Together, these findings suggest that sex differences in agonistic behavior are more nuanced than previously suggested by studies focusing primarily on offensive aggression, with males and females displaying different patterns of offensive, defensive, and avoidance behaviors during a juvenile social encounter.

Previous research has demonstrated that males generally display more overt physical aggression than females (Bell, 2018; Been et al., 2019). However, much of this research has used experimental paradigms designed to elicit aggression, including conditions involving territorial intrusion or perceived threat. (Miczek, 1979; Koolhaas et al., 2013). The present study sought to determine if sex differences in aggressive behavior would persist under social conditions in which aggression is not intentionally elicited. Under these conditions, males exhibited greater aggressive grooming and pinning, whereas females displayed more kicking and avoidance, demonstrating distinct patterns of offensive and defensive responses even when overt aggression was relatively uncommon. Importantly, composite analyses revealed a significant sex difference when offensive behaviors were considered alone, whereas no significant sex difference was detected when defensive kicking was incorporated, nor when darting was additionally included. These findings demonstrate that sex differences in individual agonistic behaviors may be obscured when offensive and defensive responses are combined into a single measure. Previous work has similarly demonstrated that sex differences in aggression are not necessarily reflected in overall aggression levels but may instead emerge in the patterns of behavioral responses expressed (Aubry et al., 2022). Together, these findings suggest that sex differences in agonistic behavior may be characterized not only by differences in the amount of aggression expressed, but also by differences in the types of behavioral responses exhibited.

Relatively few studies have directly examined the effects of VWR on aggressive behavior in rodents, and existing findings have been mixed. Ren and Tai (2023) reported that VWR reduced the frequency and duration of attacks, which included fighting, biting, chasing, and mounting, in socially isolated male mice during resident–intruder testing. Umamichi et al. (2026) found that VWR did not significantly reduce attacks in socially isolated male mice during direct interaction with an unfamiliar, age-matched conspecific, although excessive social sniffing was significantly reduced and there was a trend for exercise to reduce flight behavior. These differing findings may reflect differences in experimental conditions, including the timing and duration of social isolation and exercise, as well as the specific behaviors quantified. In the current study, VWR did not significantly affect offensive aggression, defensive aggression, or avoidance behaviors, although a trend toward reduced aggressive grooming was observed in males. However, because of the relative infrequency of aggressive behaviors during juvenile social exploration, low baseline levels of these behaviors may have limited our ability to detect effects of exercise. Consistent with this possibility, Zhao & Wang (2025) found that exercise did not significantly reduce aggression among participants with low baseline aggression, although significant effects were found in individuals with higher levels of aggression at baseline (Zhao & Wang, 2025). Additionally, behaviors were quantified during a three-minute interaction, which may have limited opportunities for aggressive responses to occur. Longer observation periods may allow for greater expression of these behaviors and improve our ability to detect differences between experimental groups. Interestingly, even this short juvenile exploration period increases circulating corticosterone in males; an effect that is attenuated by prior VWR (Greenwood et al., 2012). VWR, therefore, seems to have differential effects on behavioral and neuroendocrine responses to unthreatening social encounters. Additional studies will be needed to determine whether exercise differentially influences specific agonistic responses across social contexts.

Previous studies have reported differing effects of ovariectomy on aggressive behavior in female rats. DeBold and Miczek (1984) found that female resident rats who had been cohoused with a male rat displayed aggression toward unfamiliar female intruders following ovariectomy, specifically; attacking, bites, and aggressive posturing. In contrast, Ho et al. (2001) reported that aggression, measured using a composite score incorporating attacks, biting, on-top behavior, and piloerection, was reduced following ovariectomy, and that cyclical administration of estradiol and progesterone reinstated aggressive responding. Both studies used resident intruder paradigms, but differences in experimental variables, such as housing conditions, may have contributed to differing findings.

The present study extends this work by demonstrating that females displayed a distinct pattern of agonistic behavior during juvenile social exploration, characterized by defensive kicking and relatively little offensive aggression. Ovariectomy reduced defensive kicking, suggesting that ovarian status contributes to the expression of this female pattern of aggression. In contrast, avoidance and darting, two behaviors displayed more in females, remained unaffected by ovariectomy. These findings suggest that these avoidance behaviors are regulated through different mechanisms than defensive kicking, which is more sensitive to ovarian hormone status than other behavioral responses observed during juvenile social exploration. However, because ovariectomy removes ovarian hormone production broadly, the present findings cannot identify the contributions of individual hormones or establish the mechanisms underlying these behavioral differences.

Importantly, previous studies have demonstrated that female rats are capable of expressing offensive aggression toward unfamiliar adult conspecifics, including attacking and biting during resident–intruder testing (DeBold & Miczek, 1984; Ho et al., 2001). Thus, the limited offensive aggression observed in females in the present study may reflect the relatively nonprovocative nature of using a juvenile rather than adult social partner. Future studies directly comparing responses toward juvenile versus adult conspecifics would help determine whether the expression of offensive and defensive aggression in females depends on the social context.

Taken together, the current results demonstrate that sex differences in agonistic behavior extend beyond overall levels of aggression to distinct patterns of offensive aggression, defensive aggression, and avoidance. Males displayed greater offensive aggression, whereas females exhibited greater defensive aggression (kicking) and avoidance behavior. Ovariectomy reduced female-specific defensive aggression in females while leaving avoidance intact, while VWR did not significantly alter any of the behaviors measured. Importantly, these findings emphasize that sex differences may be obscured when distinct behavioral responses are combined into a single measure. Distinguishing offensive aggression from defensive and avoidance responses is therefore critical for understanding how sex and other biological and environmental factors influence aggressive behavior. More broadly, recognizing these behavioral distinctions may improve the interpretation of preclinical models used to investigate the underlying mechanisms of aggression and related behaviors.

## Acknowledgements

The graphical abstract was created with BioRender.

## Declaration of generative AI and AI-assisted technologies in the manuscript preparation process

During the preparation of this work, the author(s) used ChatGPT and Perplexity for clarity and brevity of wording. After using these tools, the author(s) reviewed and edited the content as needed and take full responsibility for the content of the published article.

---

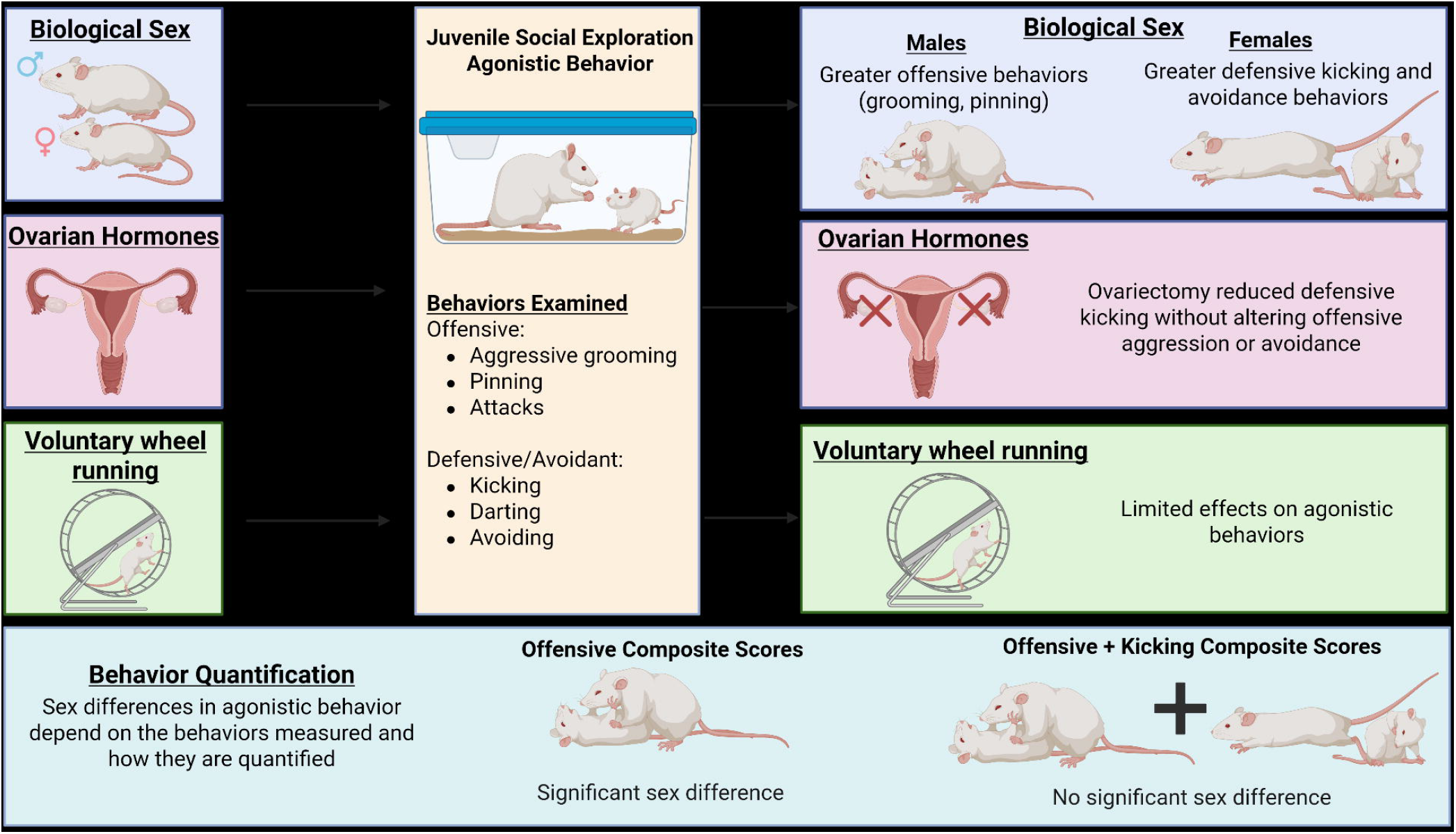

## Notes

https://zenodo.org/records/23986414?token=eyJhbGciOiJIUzUxMiJ9.eyJpZCI6ImU1NTEwZDU0LWU2NGMtNGY5NS05YzQxLWI5ZmM5MGVlYjY3MiIsImRhdGEiOnt9LCJyYW5kb20iOiI3MGJhMmJlYjA5MmNjODYxZmNlYjQyZDg2Y2U5MGU5ZSJ9.GDtfepFKm5BrY8lI5dE4D8RP_tpDaJ_7DIWRpRVSKj_xhNzc81B6pmwalY-fQo3A3aFoHcapvBSNUHhUow9Pkw

### Competing Interest Statement

The authors have declared no competing interest.

### Funding

- National Institute of Mental Health: R01MH125898

